# Interpreting blood GLUcose data with R package iglu

**DOI:** 10.1101/2020.09.28.310482

**Authors:** Steven Broll, Jacek Urbanek, David Buchanan, Elizabeth Chun, John Muschelli, Naresh M. Punjabi, Irina Gaynanova

## Abstract

Continuous Glucose Monitoring (CGM) data play an increasing role in clinical practice as they provide detailed quantification of blood glucose levels during the entire 24-hour period. The R package `iglu` implements a wide range of CGM-derived metrics for measuring glucose control and glucose variability. The package also allows to visualize CGM data using time-series and lasagna plots. A distinct advantage of `iglu` is that it comes with a point-and-click graphical user interface (GUI) which makes the package widely accessible to users regardless of their programming experience. Thus, open-source and easy to use `iglu` package will help advance CGM research and CGM data analyses. R package iglu is publicly available on CRAN and at https://github.com/irinagain/iglu.

## 1 Introduction

Continuous Glucose Monitors (CGMs) are small wearable devices that record measurements of blood glucose levels at frequent time intervals. As CGM data provide a detailed quantification of the variation in blood glucose levels, CGMs play an increasing role in clinical practice (Rodbard 2016). While multiple CGM-derived metrics to assess the quality of glycemic control and glycemic variability have been developed (Rodbard 2009*a*), their complexity and variety pose computational challenges for clinicians and researchers. While some metrics (e.g. mean) can be directly calculated from the data, others require additional pre-processing steps, such as projecting glucose measurements on equidistant time grid (e.g. CONGA, SdBDM) or the imputation of missing data.

We are aware of two existing R packages for CGM data analyses: `CGManalyzer` (Zhang et al. 2018) and `cgmanalysis` (Vigers et al. 2019). These packages are primarily designed to read and organize CGM data, rather than provide an easy-to-use interface for a comprehensive evaluation of available CGM characteristics. While their analytical utility is undeniable, a substantial number of CGM metrics summarized in Rodbard (2009*a*) is not available. Moreover, both packages require the users to have considerable programming experience, which might be a limiting factor for researchers seeking robust and accessible analytical solutions. Thus, there remains a need for open-source software that (i) computes most of CGM metrics available from the literature, and (ii) is accessible to researchers with varying levels of programming experience.

Our R package `iglu` calculates all CGM metrics summarized in Rodbard (2009*a*) in addition to several others (Bergenstal et al. 2018, Clarke & Kovatchev 2009, Danne et al. 2017), a full list of currently implemented metrics is summarized in Table 1. A comparison of functionality with `CGManalyzer` (Zhang et al. 2018) and `cgmanalysis` (Vigers et al. 2019) is in the Appendix. Additional improvements include advanced visualization with lasagna plots (Swihart et al. 2010), and provided example CGM datasets that make it easy to get started. Finally, a distinct advantage of `iglu` over existing open-source CGM software is a point-and-click graphical user interface (GUI) which makes the package accessible to users with little to no R experience.

**Table 1:**
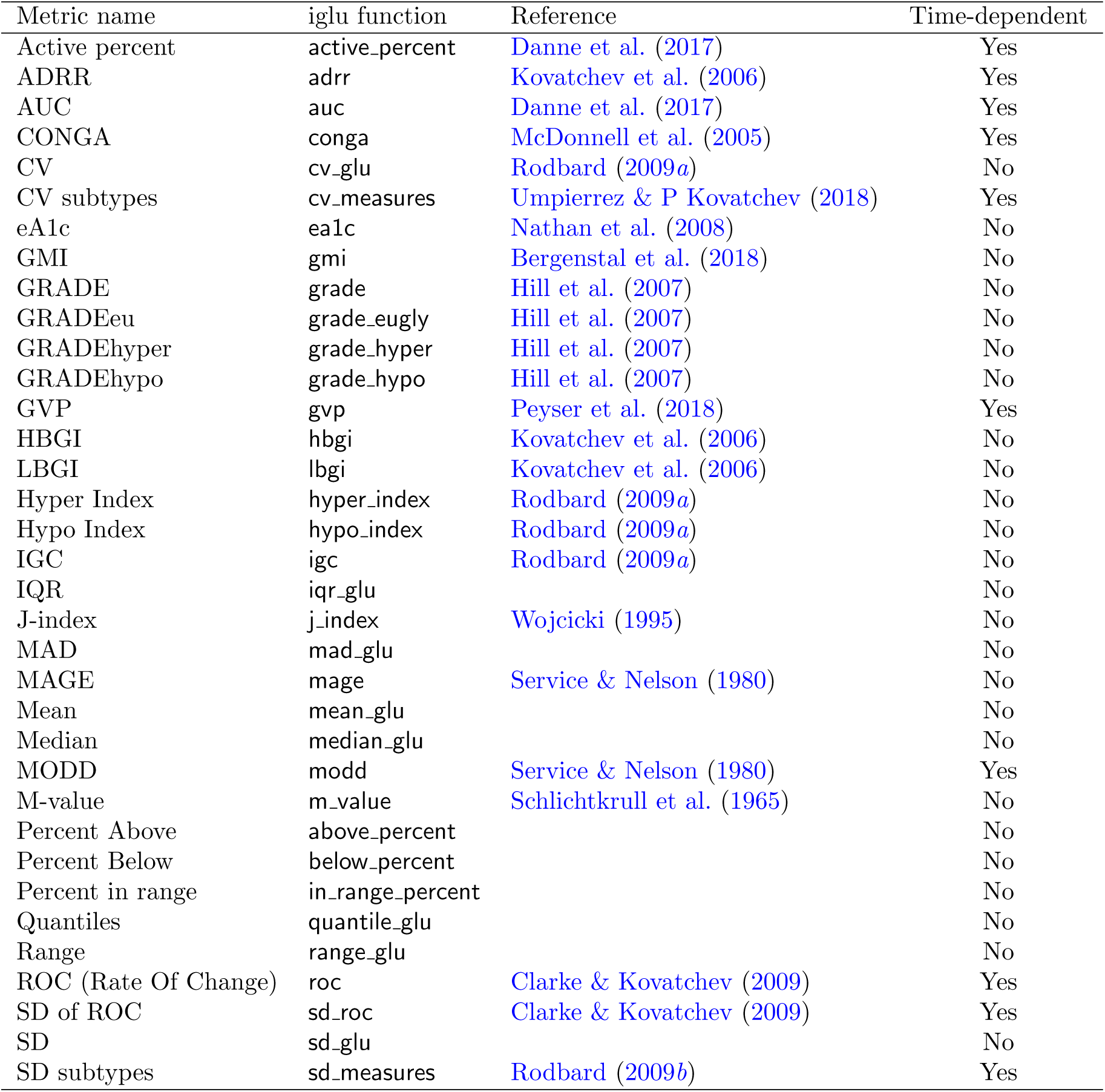
Summary of CGM metrics implemented in iglu

## 2. Features

### 2.1 Example data

The `iglu` package is designed to work with CGM data provided in the form of a data frame with three columns: id (subject identifier), time (date and time stamp) and gl (corresponding blood glucose measurement in mg/dL). The package comes with two example datasets that follow this structure. `example_data_5_subject` contains Dexcom G4 CGM measurements from subjects with Type II diabetes.

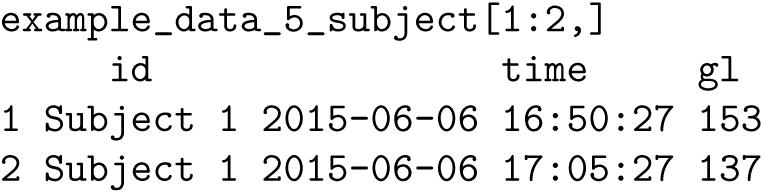

`example_data_1_subject` is a subset corresponding to one subject. These data are part of a larger study analyzed in Gaynanova et al. (2020).

### 2.2 Illustration of metrics use

Table 1 summarizes all the metrics implemented in the package, which can be divided into two categories: time-independent and time-dependent. All the functions assume that the glucose value are given in mg/dL units.

One example of a time-independent metric is Hyperglycemia index (Rodbard 2009*a*), the corresponding `iglu` function returns a single value for each subject in a tibble object (Müller & Wickham 2020). Subject id will always be printed in the id column, and metrics will be printed in the following columns.

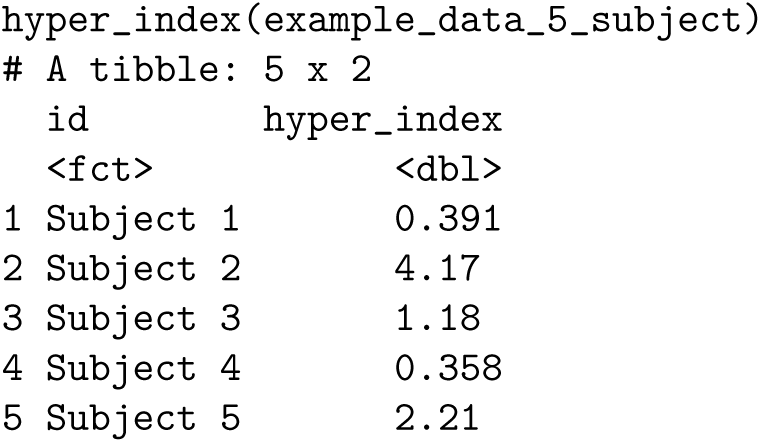

In this example, Subject 2 has the largest Hyperglycemia index, indicating the worst hyperglycemia. This is reflected in percent of times Subject 2 spends above fixed glucose targets.

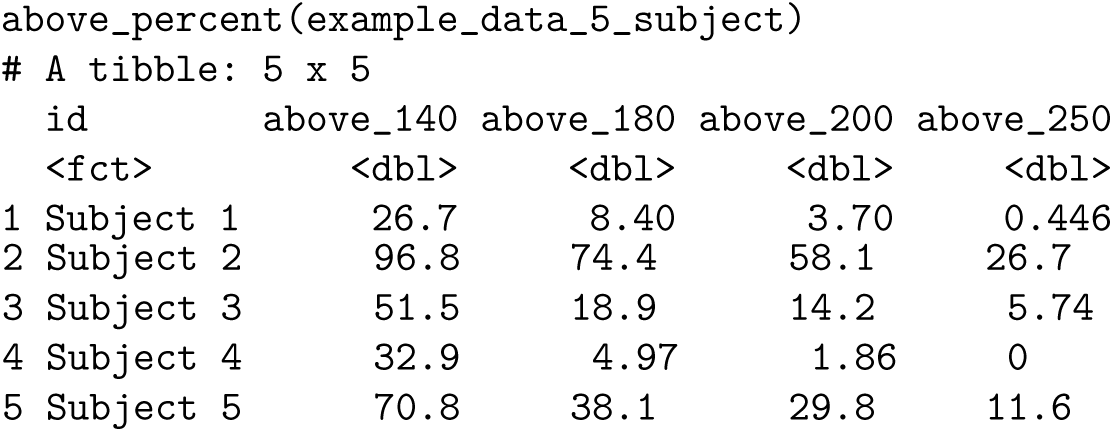

The default target values in `above_percent` can be adjusted by the user.

Examples of time-dependent metrics include measures of glycemic variability such as CONGA (McDonnell et al. 2005) and standard deviation of rate of change (Clarke & Kovatchev 2009). In the example data, the standard deviation is the highest for Subject 5 rather than for Subject 2:

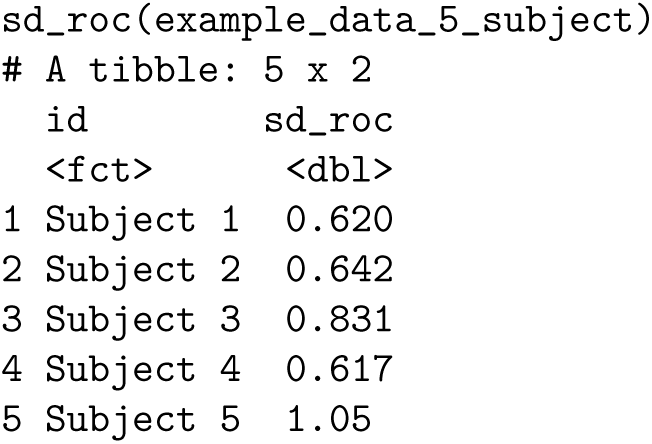

This provides an additional level of CGM data interpretation, since frequent or large glucose fluctuations may contribute to diabetes-related complications independently from chronic hyperglycemia (Suh & Kim 2015). Other metrics of glycemic variability confirm the high fluctuations in Subject 5, with all but one subtypes of standard deviation being the largest for Subject 5 (Rodbard 2009*b*):

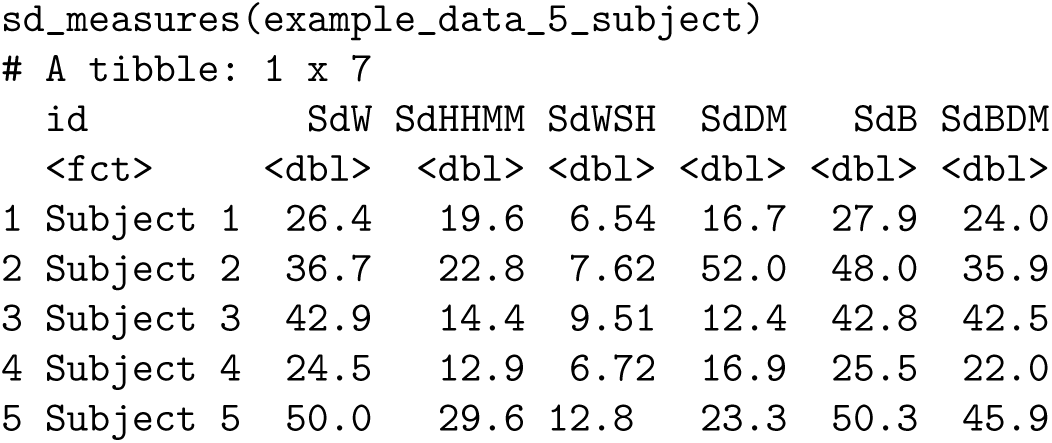

The calculations of these variability metrics require evenly spaced glucose measurements across time; however this is not always the case in practice due to missing values. In order to create an evenly spaced grid of glucose measurements, `iglu` provides the function `CGMS2DayByDay`. This function is automatically called for metrics requiring the evenly spaced grid, however the user can also access the function directly. The function works on a single subject data, and has three outputs.

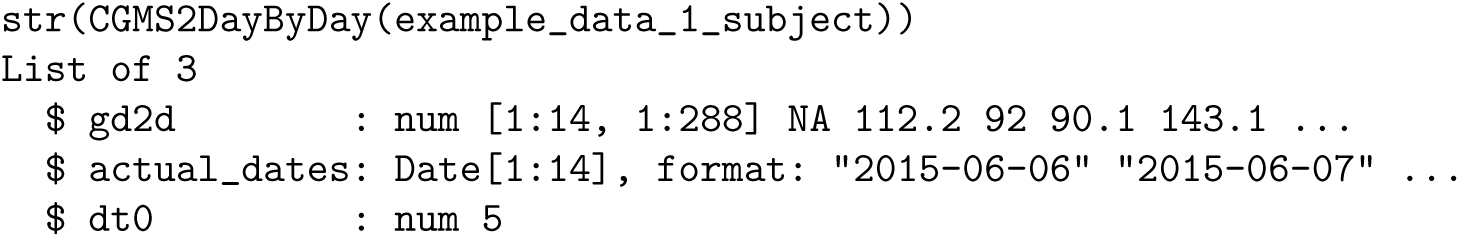

The first part of the output, `gd2d`, is the interpolated grid of values. Each row correspond to one day of measurements, and the columns correspond to equi-distant time grid covering 24 hour time span. The grid is chosen to match the frequency of the sensor (5 minutes in this example leading to (24 ∗ 60)*/*5 = 288 columns), which is returned as `dt0`. The returned `actual_dates` allow to map the rows in `gd2d` back to original dates. The achieved alignment of glucose measurement times across the days enables both the calculation of corresponding metrics, and the creation of lasagna plots (Section 2.3).

### 2.3 Visualizations

The iglu package has several visualization capabilities, which are summarized in Table 2. The main function is `plot_glu`, which by default provides time series plot for each subject. Figure 1 illustrates the output on example data with the horizontal red lines indicating user-specified target range. The visual inspection of the plots confirm the conclusions of Section 2.2, most of the measurements for Subject 2 are above 140 mg/dL, however the variability is larger for Subject 5.

**Table 2:** Summary of iglu visualization capabilities

| Function call | Visualization description | Main parameters |
| --- | --- | --- |
| `plot_glu` | Multiple plot types: time series and lasagna | `plottype`, `lasagnatype` |
| `plot_lasagna` | Lasagna plot of glucose values for multiple subjects | `datatype`, `lasagnatype` |
| `plot_lasagna_1subject` | Lasagna plot of glucose values for a single subject | `lasagnatype` |
| `plot_roc` | Time series of glucose values colored by rate of change (ROC) | `subjects`, `timelag` |
| `hist_roc` | Histogram of rate of change (ROC) values | `subjects`, `timelag` |

**Figure 1:**
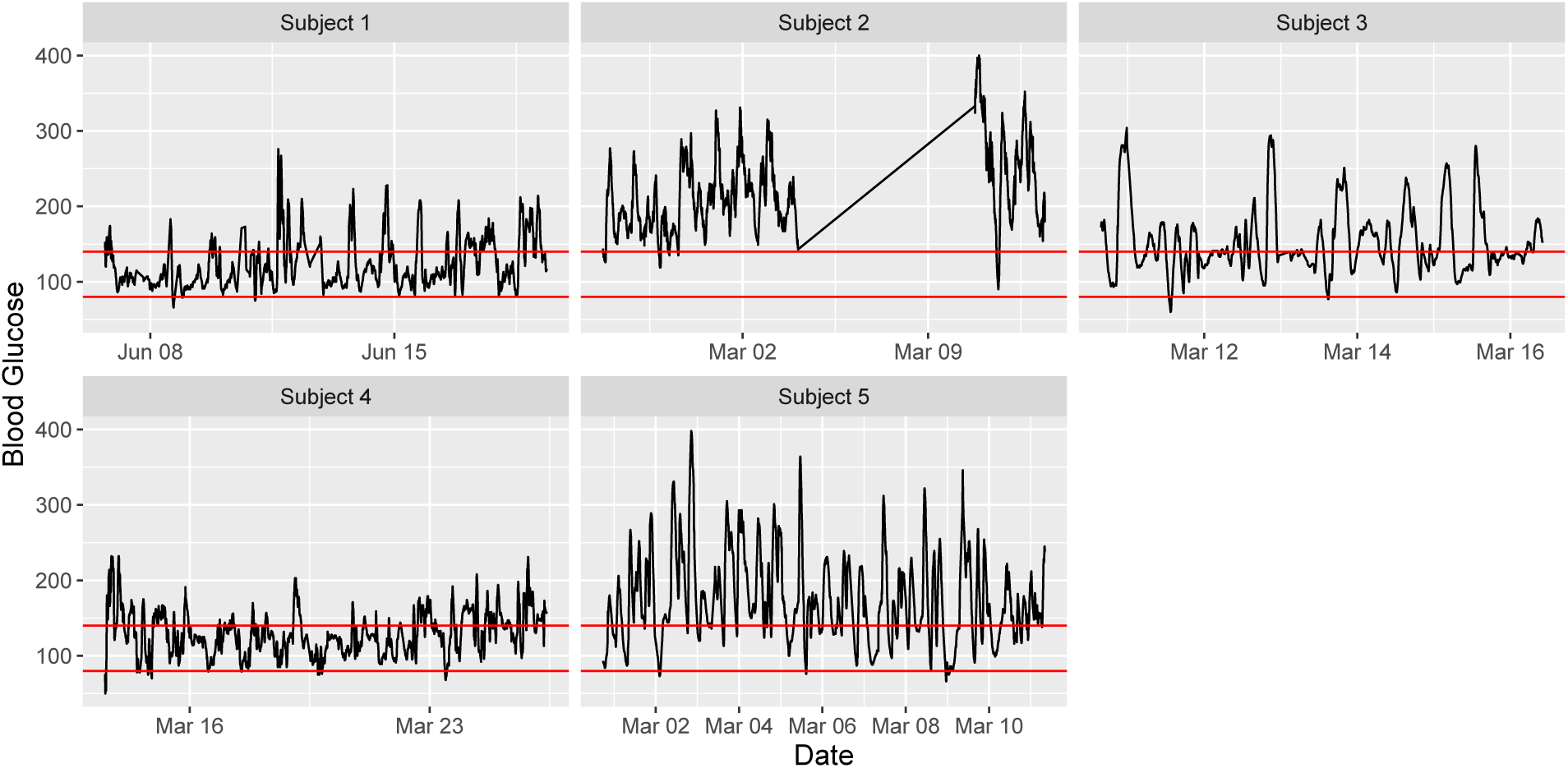
Time series plots for five subjects with selected target range [80, 140] mg/dL. The linear trend in the middle of Subject 2 plot is an artifact of missing glucose values for that time range. While visible in the time-series plot, this artifact does not affect the metrics calculation.

Another visualization type is provided via lasagna plots (Swihart et al. 2010), which use color grid rather than the number scale to visualize trends in data over time. The lasagna plots in `iglu` can be single-subject or multi-subject. The single-subject lasagna plot has rows corresponding to each day of measurements with color grid indicating glucose values (Figure 2 **A**). An optional within-time sorting across days allows to investigate average glucose patterns as a function of 24 hour time period (Figure 2 **B**). The multi-subject lasagna plot has rows corresponding to subjects, with color grid indicating glucose values across the whole time domain, or average glucose values across days. The highest glucose values are displayed in red, whereas the lowest are displayed in blue. Thus, the numerical glucose values are mapped to color using the gradient from blue to red (Figure 2). The functions allow user to modify the default gradient scale. Figure 2 **C** displays customized multi-subject lasagna plot for example data that displays average glucose values across days for each subject, this plot is produced by the following call.

**Figure 2:**
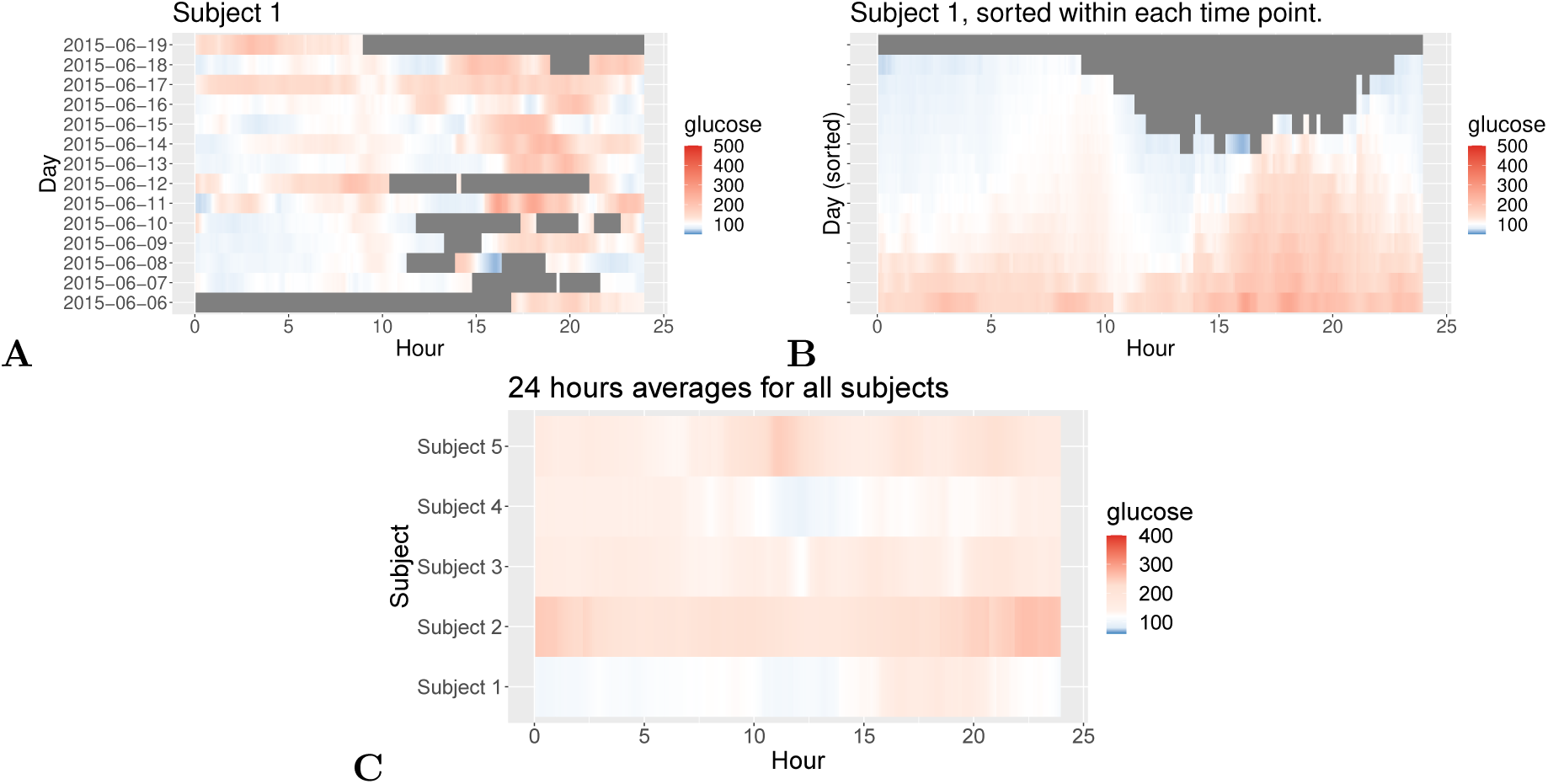
**(A)** unsorted and **(B)** time-sorted lasagna plot for Subject 1; **(C)** unsorted customized multi-subject lasagna plot based on average values across days

~~~
plot_lasagna(example_data_5_subject, datatype = “average”,
                      midpoint = 140, limits = c(60, 400))
~~~

The `midpoint` specifies the glucose value (in mg/dL) at which the color transitions from blue to red (the default is 105 mg/dL), whereas the `limits` specify the range (the default is [50, 500] mg/dL). From Figure 2 one can for example infer that the glucose values for Subject 1 tend to be the highest in late afternoon (≈ 15:00 - 20:00). One can also infer that Subject 1 tends to have the lowest glucose values during night time hours (0:00 - 6:00) compared to other four subjects.

In addition to visualizing absolute glucose values, iglu also allows to visualize local changes in glucose variability as measured by rate of change (Clarke & Kovatchev 2009). There are two types of visualizations associated with rate of change. The first is a time series plot of glucose values where each point is colored by the rate of change at that given time. Points colored in white have a stable rate of change, meaning the glucose is neither significantly increasing nor decreasing at that time point. Points colored red or blue represent times at which the glucose is significantly rising or falling, respectively. Thus colored points represent times of glucose variability, while white points represent glucose stability. Figure 3 **A** shows a side by side comparison of rate of change time-series plots for two subjects. Subject 1 shows significantly less glucose variability than Subject 5. The function call to produce this plot is as follows.

**Figure 3:**
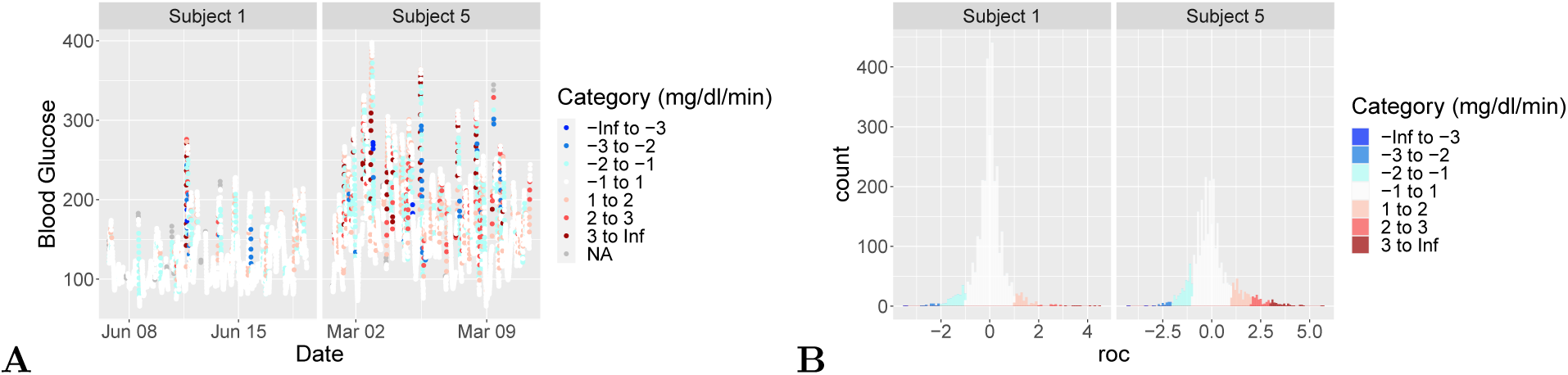
Rate of change visualizations: **(A)** time-series and **(B)** histogram plots of rate of change for two selected subjects from example dataset.

~~~
plot_roc(example_data_5_subject, subjects = c(”Subject 1”, “Subject 5”))
~~~

Figure 3 **B** shows a side by side comparison of rate of change histogram plots for the same subjects. Once again, the colors show in what direction and how quickly the glucose is changing. The histogram plots allow to immediately assess the variation in rate of change. Extreme values on either end of the histogram indicate very rapid rises or drops in glucose - a high degree of local variability. In Figure 3, Subject 1 once again shows lower glucose variability by having a narrower histogram with most values falling between −2 mg/dl/min and 2 mg/dl/min. Subject 5 has a shorter, more widely distributed histogram indicating greater glucose variability. The function call to produce this plot is as follows.

~~~
hist_roc(example_data_5_subject, subjects = c(”Subject 1”, “Subject 5”))
~~~

### 2.4 Relationship between metrics

To illustrate the relationships between different metrics and their intepretation, we calculate all metrics for example data of 5 subjects (Section 2.1). Figure 4 shows heatmap of resulting metrics (centered and scaled across all subjects to aid visualization) created using R package `pheatmap` (Kolde 2019). The hierarchical clustering of glucose metrics results in six meaningful group with the following interpretation (from top to bottom): (1) in range metrics; (2) hypoglycemia metrics; (3) hyperglycemia metrics; (4) a mixture of variability and hyperglycemia metrics; (5) CVsd (standard deviation of CV, coefficient of variation, across days); (6) glucose variability metrics. Interestingly, while CVsd is a measure of glucose variability, it behaves quite differently from other variability metrics in these 5 subjects. The hierarchical clustering of subjects confirms our previous observations that Subjects 2 and 5 have worse glucose control compared to Subjects 1, 3 and 5. Furthermore, it confirms that Subject 2 has the worst hyperglycemia (highest values for metrics in group (2)), whereas Subject 5 has the highest glucose variability (highest values for metrics in group (6)).

**Figure 4:**
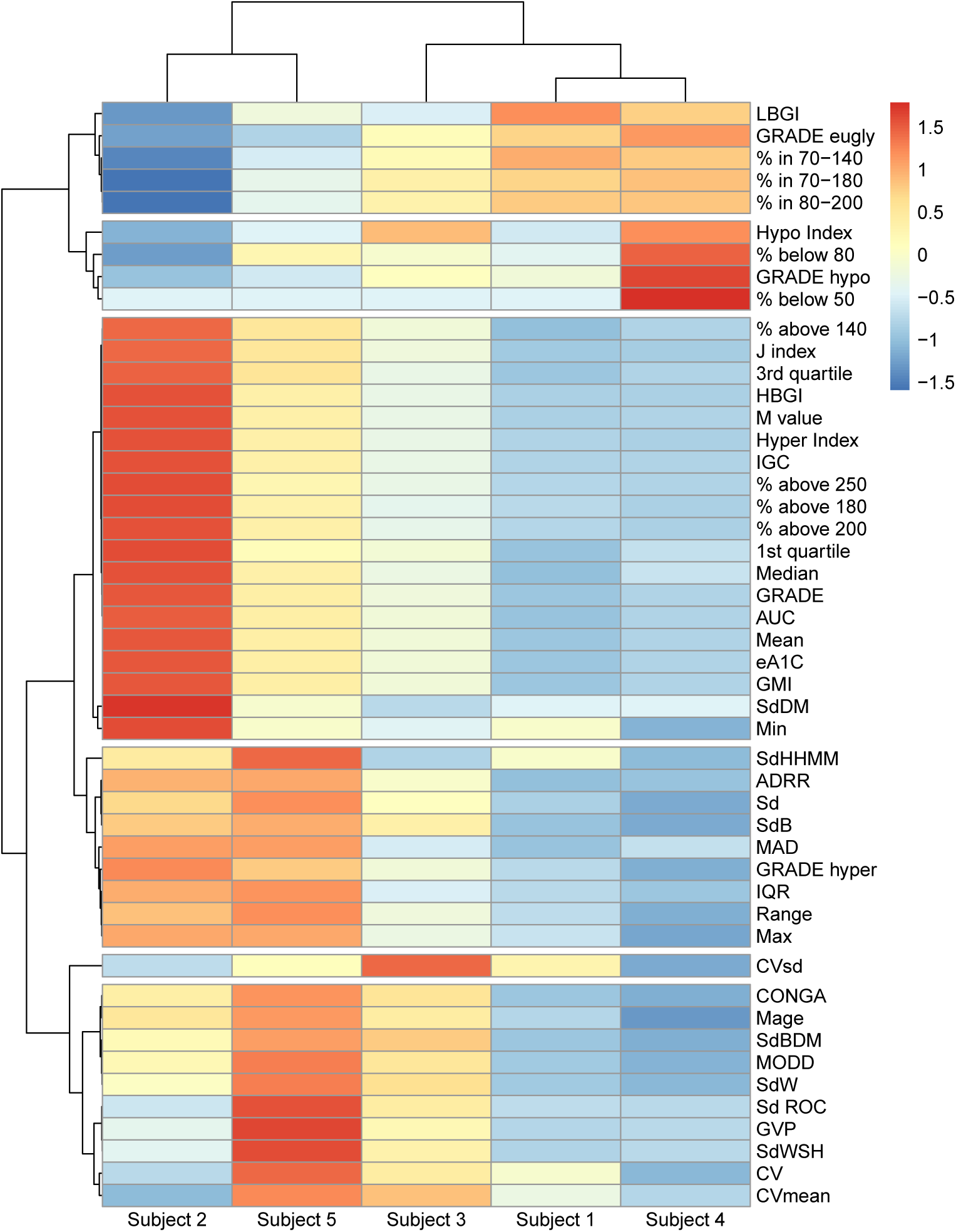
Heatmap of all metrics calculated using `iglu` for 5 subjects with Type II diabetes, hierarchical clustering is performed on centered and scaled metric values using distance correlation and complete linkage. The cluster tree for metrics is cut at 6 groups, which can be interpreted as follows (from top to bottom): (1) in range metrics; (2) hypoglycemia metrics; (3) hyperglycemia metrics; (4) a mixture of variability and hyperglycemia metrics; (5) CVsd (standard deviation of CV, coefficient of variation, across days); (6) glucose variability metrics. The heatmap supports that Subjects 2 and 5 have worse glucose control than then other subjects, with Subject 2 having the worst hypoglycemia and Subject 5 the highest variability.

### 2.5 GUI via shiny application

The iglu package comes with a shiny application (Chang et al. 2020), which provides a point-and-click graphical user interface (GUI) for all metric calculations and visualizations. The interface can be accessed from R console by calling

~~~
iglu::iglu_shiny()
~~~

or directly at https://irinagain.shinyapps.io/shinyiglu/. The users can load their CGM data in .csv format, and export metrics output to the user’s clipboard or to .csv, .xlsx, or .pdf files (Figure 5 A-B). Figure 5 C shows an example of shiny interface for creating customized visualization plots based on user-loaded data.

**Figure 5:**
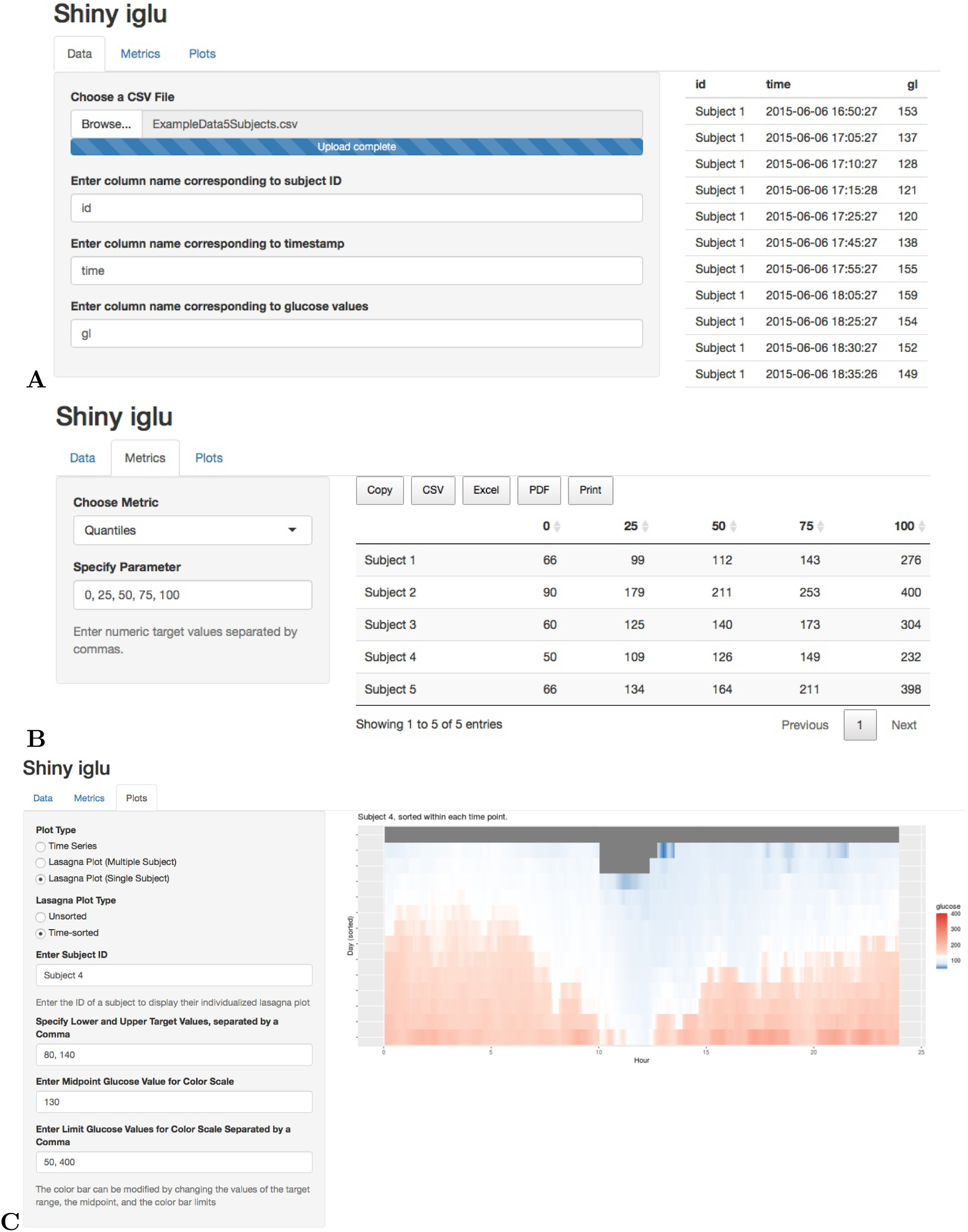
Shiny interface for **(A)** loading CGM data in .csv format; **(B)** calculating user-specified quantiles for each subject; **(C)** creating customized lasagna plot for the selected subject.

## 3 Discussion and conclusion

The `iglu` package is designed to simplify computations of CGM-derived glucose metrics, and assist in CGM data visualization. The current version includes all of the metrics summarized in Rodbard (2009a) as well as many others (see Table 1). New metrics will be incorporated into the future versions as they develop. More details on the package functionality are provided in the package vignette available at https://irinagain.github.io/iglu/.

While there are existing open-source R packages for CGM data analyses (Vigers et al. 2019, Zhang et al. 2018), these packages focus more on CGM data reading than exhaustive metric implementation, and require programming experience. Instead, `iglu` focuses on comprehensive implementation of available CGM metrics and ease of use via accompanying GUI application. All data loading, parameter selection, metric calculations and visualizations are available via point-and-click graphical user interface. This makes `iglu` accessible to a wide range of users, which coupled with free and open-source nature of `iglu` will help advance CGM research and CGM data analyses.

## Acknowledgements

The authors are thankful to Marielle Hicban, Mary Martin, Nhan Nguyen, Pratik Patel and John Schwenck for assisting in writing several metric functions in version 2.0.0 of iglu package. This work was supported by the National Institutes of Health [HL11716, HL146709]; and an agreement from the Johns Hopkins University [80045538 to I.G.].

## A Comparison of iglu functionality with existing packages

Table 3 compares implemented CGM metrics from `iglu` with `CGManalyzer` (Zhang et al. 2018) and `cgmanalysis` (Vigers et al. 2019).

**Table 3:** Comparison of R packages on CGM metrics

| Metric name | CGManalyzer | cgmanalysis | iglu |
| --- | --- | --- | --- |
| Summary statistics | ✓ | ✓ | ✓ |
| Active percent | × | ✓ | ✓ |
| ADRR | × | × | ✓ |
| AUC | × | ✓ | ✓ |
| CONGA | ✓ | ✓ | ✓ |
| CV subtypes | × | × | ✓ |
| eA1c | × | ✓ | ✓ |
| GMI | × | ✓ | ✓ |
| GRADE | × | × | ✓ |
| GVP | × | × | ✓ |
| HBGI/LBGI | × | ✓ | ✓ |
| IGC | × | × | ✓ |
| J-index | × | ✓ | ✓ |
| MAGE | × | ✓ | ✓ |
| MODD | ✓ | ✓ | ✓ |
| M-value | × | × | ✓ |
| Percent in range | × | ✓ | ✓ |
| ROC (Rate of Change) | × | × | ✓ |
| SD subtypes | × | × | ✓ |
| GUI for calculation | × | × | ✓ |

